# The acetyltransferase Eco1 elicits cohesin dimerization during S phase

**DOI:** 10.1101/2020.02.07.938530

**Authors:** Di Shi, Shuaijun Zhao, Mei-Qing Zuo, Jingjing Zhang, Wenya Hou, Meng-Qiu Dong, Qinhong Cao, Huiqiang Lou

## Abstract

Sister chromatid cohesion is established by Eco1 in S phase. Nevertheless, the exact consequence of Eco1-catalyzed acetylation is unknown, and the cohesive state remains highly controversial. Here we show that self-interactions of cohesin subunits Scc1/Rad21 and Scc3 occur in a DNA replication-coupled manner in both yeast and human. Through cross-linking mass spectrometry and VivosX analysis of purified cohesin, we show that a subpopulation of cohesin may exist as dimers. Importantly, cohesin-cohesin interaction becomes significantly compromised when Eco1 is depleted. On the other hand, deleting either deacetylase Hos1 or Eco1 antagonist Wpl1/Rad61 results in an increase (e.g., from ∼20% to 40%) of cohesin dimers. These findings suggest that cohesin dimerization is controlled by common mechanisms as the cohesion cycle, thus providing an additional layer of regulation for cohesin to execute various functions such as sister chromatid cohesion, DNA repair, gene expression, chromatin looping and high-order organization.

**Author Summary:** Cohesin is a ring that tethers sister chromatids since their synthesis during S phase till their separation in anaphase. According to the single-ring model, one ring holds twin sisters. Here we show a conserved cohesin-cohesin interaction from yeast to human. A subpopulation of cohesin is dimerized concomitantly with DNA replication. Cohesin dimerization is dependent on the acetyltransferase Eco1 and counteracted by the anti-establishment factor Wpl1 and deacetylase Hos1. Approximately 20% of cellular cohesin complexes are measured to be dimers, close to the level of Smc3 acetylation by Eco1 in vivo. These findings provide evidence to support the double-ring model in sister chromatid cohesion.

## Introduction

Cohesin is a tripartite ring that controls many, if not all, aspects of chromosome function including sister chromatid cohesion, chromosome segregation, chromosome condensation, chromosome organization, DNA replication, DNA repair, DNA recombination and gene expression (1-5). The ring consists of a V-shaped heterodimeric SMC proteins Smc1 and Smc3, and an α-kleisin subunit Scc1/Rad21 bridging their ABC ATPase head domains (6, 7). The fourth subunit, Scc3 (SA1 or SA2 in mammalian cells), binds to the ring through Scc1 (8). The stoichiometry of these subunits is 1:1:1:1(9-11). Besides such a single-ring model (12-18), higher-order oligomeric cohesin conformations have also been proposed based upon the unusual genetic properties and physical self-interactions of cohesin subunits in yeast (19) and human cells (20), respectively. Therefore, it remains highly debatable whether cohesin functions as single-rings, double-rings or clusters (6, 21-23).

As an essential process mediated by cohesin, sister chromatid cohesion is achieved in two steps, cohesin loading and cohesion establishment (24, 25). Cohesin is loaded onto chromosomes with a low affinity in G1 phase (26). Following DNA replication, cohesion is established wherein it tightly holds twin chromatids together to resist the pulling force of the spindle before segregation (27, 28). Eco1 plays an essential role in the establishment of cohesion (29-32). It acetylates two conserved lysine residues on Smc3 (K112 and K113 in *Saccharomyces cerevisiae*) (33-35). Smc3 acetylation appears to counteract an ‘‘anti-establishment’’ activity of Wpl1/Rad61 (36). Wpl1 ablation restores viability and improves sister chromatid cohesion in the absence of Eco1 or Smc3 acetylation(33, 37, 38). Nevertheless, the exact consequences of Smc3 acetylation remain unknown either.

## Results and discussion

### Self-interactions of cohesin subunits in yeast cells

Self-interactions of cohesin subunits have not been detected in yeast (9). In human cells, contradictory observations have been reported (20, 39). To clarify this, we first performed immunoprecipitation (IP) experiments using a similar dual-tag strategy. An ectopic copy of *SCC1* (pRS-*SCC1*, Table S1) under the control of its native promoter was introduced into a haploid yeast strain. The two *SCC1* alleles were labeled with a pair of orthogonal epitopes (GFP/FLAG or EPEA/FLAG), which has been well demonstrated to be orthogonal to each other in this and previous studies (Figure 1)(40, 41). When the epitopes were inserted at the C-termini of both copies (Scc1-HA-EPEA/Scc1-5FLAG), after EPEA-IP, no Scc1-Scc1 interaction was virtually detectable (data not shown), in agreement with a previous study from the Nasmyth’s group in yeast (9). Epitope tagging may occasionally cause unexpected interference on the structure and function of proteins, which turns out to be true for many cohesin subunits (20, 42). To test this possibility, we switched 5FLAG from the C-terminus to the N-terminus of Scc1. Although EPEA-IP was performed in the exact same procedure, such a change led to the positive interaction between Scc1-HA-EPEA and 5FLAG-Scc1 (Figure 1A, lane 6). Because EPEA can only be applied at the C-terminus, we then labeled both Scc1 copies at their N-termini with another orthogonal epitope pair, GFP/FLAG. A similar intermolecular interaction was observed in GBP-IP (Figure 1B, lane 6). These results are consistent with the observations in human cells (20), indicating that the self-interaction of Scc1 might be conserved.

**Figure 1.**
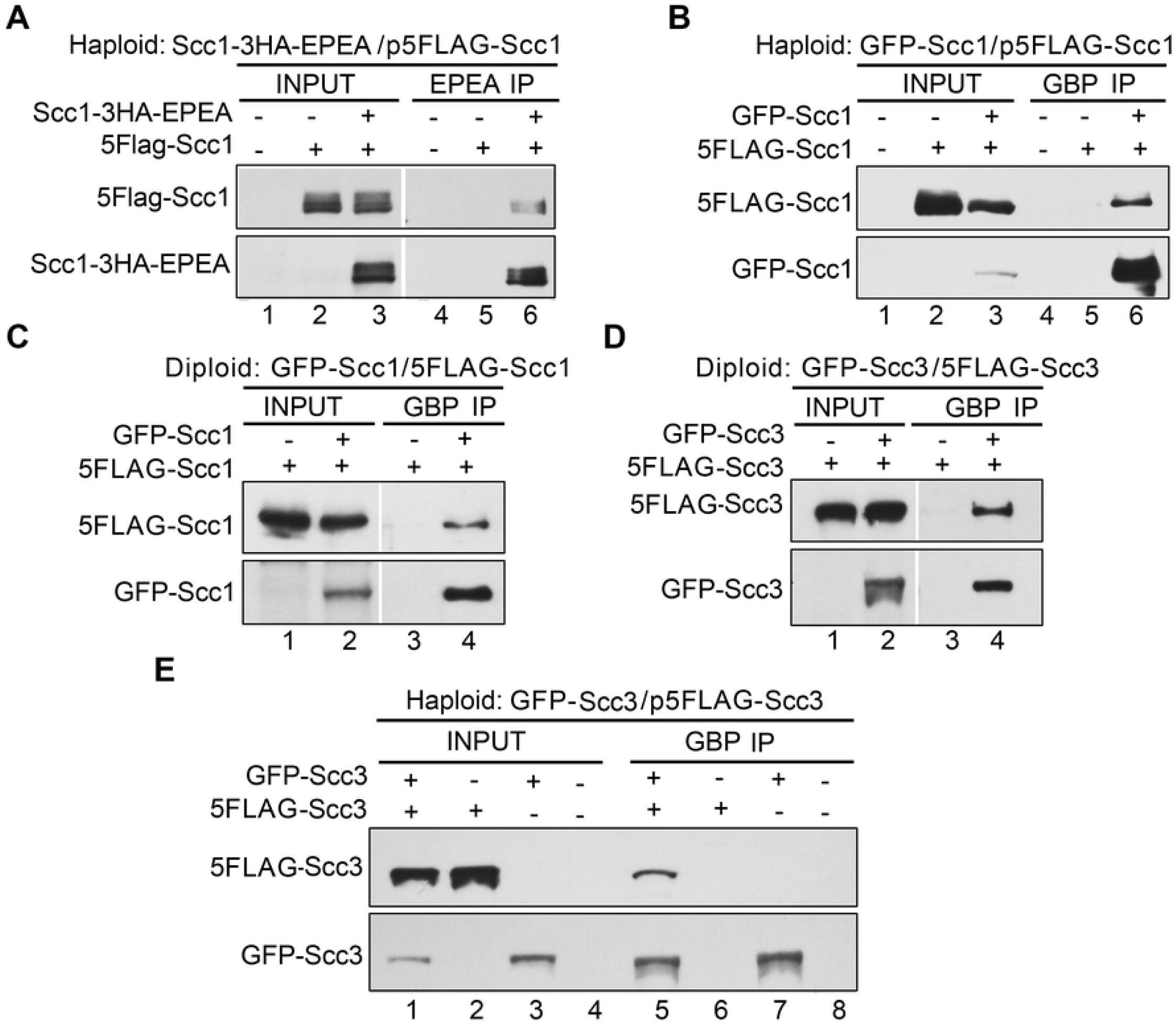
Intermolecular interactions of cohesin subunits in yeast. (A) Self-interaction of Scc1 in the overexpression condition. An extra copy of 5FLAG-SCC1 (Table S1) was introduced into the strain (Table S2, YSD24) carrying a C-terminal 3HA-EPEA-tagged *SCC1* at the genomic locus. The lysates (input) were prepared from the cells collected at the exponential phase. Scc1-3HA-EPEA was precipitated via a C-tag affinity matrix. The precipitated proteins were detected via immunoblots against FLAG and HA antibodies, respectively. (B) Self-interaction of Scc1 detected via another pair of orthogonal epitopes. A plasmid expressing an extra copy of 5FLAG-*SCC1* was introduced into the haploid yeast strain carrying an N-terminal GFP-tagged *SCC1* at the genomic locus (Table S2, YSD03). GFP-Scc1 was precipitated via GBP beads. The precipitates were analyzed via IB against FLAG and GFP antibodies, respectively. (C) Self-interaction of Scc1 in the physiological protein levels. The two endogenous *SCC1* alleles in yeast diploid cells were labeled at their N-termini with GFP and 5FLAG, respectively (Table S2, YSD107). The lysates were prepared from the cells collected at the exponential phase. GBP-IP and IB were performed as above. (D, E) Self-interaction of Scc3 in the physiological (D) or overexpression (E) condition. A diploid yeast strain (Table S2, YSD109) carrying the two endogenous *SCC3* alleles with N-terminal GFP and 5FLAG tags was used in (D). A haploid strain (Table S2, YSD61) carrying an endogenous N-terminal GFP-tagged *SCC3* and an ectopic copy of 5FLAG-*SCC3* was used in (E). GBP-IP and IB were basically performed as above.

The discrepancy could be explained if Scc1 oligomerization is interrupted by C-terminal tagging. However, it is also possible that self-interaction might be artificially caused by overexpression of cohesin subunits (22). To test the latter possibility, we next labeled two endogenous *SCC1* alleles with the same pair of orthogonal epitope tags (i.e., GFP/FLAG) at their genomic loci in the diploid yeast cells. Under the physiological protein levels, Scc1-Scc1 interaction was apparent as well (Figure 1C, lane 4), consistent with a very recent study in mouse embryonic stem cells (mESCs) (43). Intriguingly, self-interaction was also observed for the fourth cohesin subunit, Scc3, under endogenous and overexpression conditions in diploid (Figure 1D, lane 4) and haploid (Figure 1E, lane 5) cells, respectively. Collectively, these data suggest that cohesin is able to form dimers or oligomers. The failure to detect the oligomeric cohesin is due to inappropriate epitope tagging and/or other experimental conditions.

### Isolation and crosslinking analysis of the cohesin complexes

To corroborate and characterize the different cohesin species, we next purified the complexes from yeast cells containing two copies of Scc1 with small (5FLAG) and large (GST) epitopes, respectively. This allowed the simultaneous detection of the two Scc1 copies in a single gel by probing with anti-Scc1 antibodies. The lysates were first subjected to anti-FLAG affinity purification and FLAG peptide elution. The eluates were then run on a 10-30% glycerol sedimentation/velocity gradient. After centrifugation, fractions (0.5 ml each, labeled from top to bottom) were collected. After separation by SDS-PAGE, immunoblots revealed the co-purification of Smc3 together with Scc1, suggesting successful isolation of the complex rather than an individual Scc1 subunit (Figure 2A). The peak of the purified complex (fractions 6-9) contained few GST-Scc1 (i.e., the second copy of Scc1), sedimenting close to the 669 kDa standard (fraction 8). The theoretical molecular weight of the single-ring four-subunit cohesin complex is 478 kDa. The relative broad distribution of the cohesin complexes in the glycerol gradient might be due to the co-purification of the additional factors like Pds5 and Wpl1. However, the cohesin species containing the second Scc1 copy (GST-Scc1) were clearly detected, which sedimented much faster than 669 kDa, peaking around fraction 13. These data corroborate the existence of cohesin dimers/oligomers in vivo.

**Figure 2.**
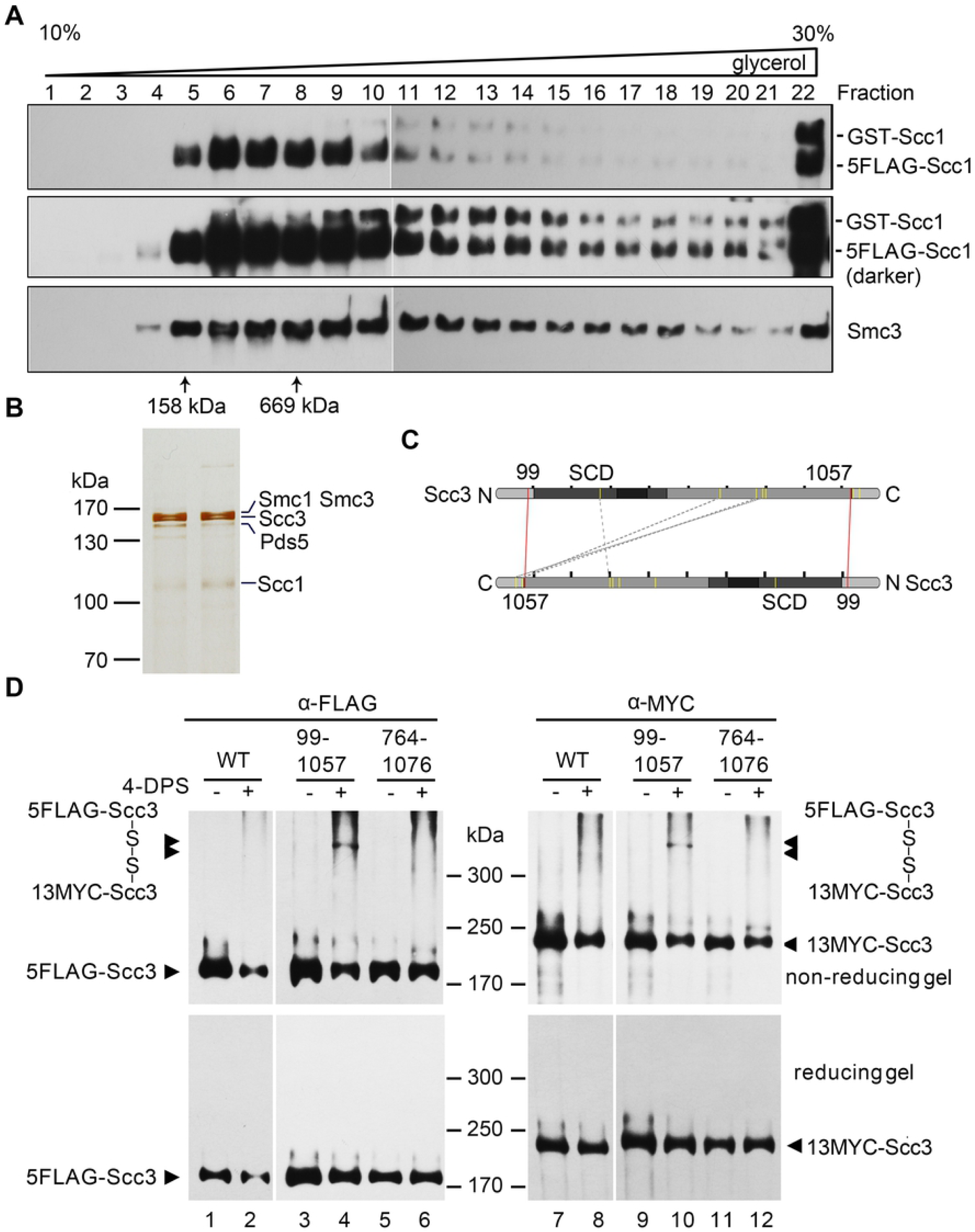
Purification and crosslinking of the cohesin complexes. (A, B) Purification of the native cohesin complexes. The cohesin complexes were isolated from the yeast cells expressing *p5FLAG-SCC1* cells via one step affinity purification (i.e. anti-FLAG M2 column chromatography and FLAG peptide elution) followed by 10-30% glycerol density gradient centrifugation. The sample was divided into 24 fractions (0.5 ml each). After separation by SDS-PAGE, they were analyzed by IB with the indicated antibodies (A) or silver staining (B). The sedimentation of standard proteins (158 and 669 kDa) is indicated by an arrow. The band of each subunit was validated by MS as well. (C) An Scc3-Scc3 connectivity map of the high-confidence DSS cross-links detected in cross-linking mass spectrometry (CXMS). The purified Scc3-containing complexes were cross-linked by DSS prior to trypsin digestion and LCMS/MS as described in Experimental Procedures. The crosslinked amino acids were identified using the pLink search engines and labeled by a dashed grey line. The pair of amino acids validated by the following cysteine substitution and in vivo crosslinking are labeled by a red line. (D) Cysteine substitution of K99 and K1057 at the putative Scc3-Scc3 interface supports the in vivo crosslinking. Cys-screening of the putative pairs near the intermolecular interface of Scc3. The pair of the indicated amino acid residues (e.g., K99/K1057; K764/K1076) in two copies of Scc3 were substituted by cysteine. WT or cysteine-substituted mutant cells were grown and treated with 180 μM 4-DPS (+) or DMSO (-) before harvest. The proteins were extracted and analyzed by non-reducing (top panel) or reducing (bottom panel) SDS-PAGE followed by anti-FLAG and anti-MYC IBs. The monomeric and dimeric Scc3 were indicated by single and double arrows, respectively.

Next, we omitted the GST tag which may cause artificial dimerization. The cohesin complexes at the endogenous level were isolated through 5FLAG-Scc1 or 5FLAG-Scc3. Silver staining showed that the cohesin complex is purified to a nearly homogeneous level (Figure 2B). Besides all four cohesin subunits, the cellular cohesins often contained other components like Pds5 as validated by liquid chromatography-mass spectrometry (LC-MS). To determine how cohesin interacts with itself, we then performed cross-linking MS (CXMS) of the purified cohesin complexes. The representative cross-linked amino acids mapped to Scc3 were shown in Figure 2C. Although it is challenging to distinguish between intramolecular and intermolecular interfaces of a homo-oligomer, we supposed that the pairs of cross-linked residues apart from each other in the available three-dimensional structure of Scc3 fragment likely represent the intermolecular interface.

To test this, we then substituted these putative pairs by cysteine substitution for the VivosX (in <u>vivo</u> di<u>s</u>ulfide <u>cross</u>linking) assay (44). If the two amino acids replaced by cysteine were close enough, a disulfide bond would be introduced by the permeable thiol-specific oxidizing agent 4,4’-dipyridyl disulfide (4-DPS). To simplify the screening and detection, two Scc3 alleles with a pair of tags (5FLAG-Scc3/13MYC-Scc3) were expressed in yeast cells. In WT, Scc3 (either 5FLAG-Scc3 or 13MYC-Scc3) migrated as a monomer (less than 200 kDa) with or without 4-DPS treatment in non-reducing SDS-PAGE (Figure 2D, lanes 1, 2, 7, 8, top panel). Among all mutated amino acid pairs, a portion of the Scc3-Scc3 crosslinking adducts was only detectable in the Scc3-K99C/Scc3-K1057C pair after 4-DPS treatment (Figure 2D, compare lanes 3-4, 9-10). They migrated more slowly than 300 kDa, close to the expected molecular weight of dimeric Scc3 (∼287 kDa). Importantly, the same band was able to be probed by both anti-FLAG (lanes 4, top panel) and anti-MYC (lanes 10, top panel), confirming that it is a dimeric/oligomeric Scc3 complex. Moreover, the band was abolished in the reducing gel (Figure 2D, bottom panel), further validating that it is formed by disulfide cross-linking of the introduced cysteine pair. Putting together, these results suggest the existence of the Scc3-Scc3 dimer in vivo. Given the distal location of K99 and K1057 at the unstructured N- and C-termini of Scc3, these results also implicate that the twin Scc3 molecules might bind each other in an antiparallel manner to mediate a double-ring form of the cohesin complex.

Notably, through bi-fluorescent complementation assays, Zhang et al have shown a similar antiparallel orientation of Scc1-Scc1 in human cells (20). However, they failed to detect intermolecular interaction of Scc3 subunit, probably due to two Scc3 homologs (SA1 and SA2) in mammals or other experimental conditions. According to the LC-MS quantification during G1, G2 and M phases, the Peters’ group concluded that the stoichiometry of cohesin complexes remains to be 1:1:1:1 (monomer) or 2:2:2:2 (dimer) (45). This finding supports the cohesin dimerization identified here and also recently in mESCs (43).

### Replication-coupled Smc3-Smc3 interaction in human cells

Since the cohesin status is cell-cycle regulated(46), we wanted to know whether cohesin dimerization is similarly controlled. To test this, we applied a proximity ligation assay (PLA) to visualize the cohesin-cohesin interaction in human cells (47). 5FLAG-Smc3 and 13MYC-Smc3 were introduced into HeLa cells. Cells were grown and arrested in G1 (0 h) by double thymidine block, before release into the fresh media containing EdU for 2, 4, 6, 8, 10 h. Two Smc3 copies were probed by mouse anti-FLAG and rabbit anti-MYC antibodies, respectively. If two Smc3 proteins are in proximity, their secondary antibodies conjugated to DNA oligonucleotides will bring together another pair of oligonucleotides, which is subsequently ligated and circulated by DNA ligase. The circulated DNA was amplified by rolling circle amplification and finally detected by fluorescence in situ hybridization (FISH). In G1 phase, few fluorescence signals were observed (Figure 2A), excluding the possible false positives presumably due to the high sensitivity of the PLA method and/or Smc3 overexpression. Post G1 release, PLA signals appeared in 2 h and peaked around 6 h (Figures 3A and 3B). These results corroborate the cohesin-cohesin interaction originally discovered by Pati’s group in human cells (20). More importantly, these data also demonstrate that cohesin oligomerization does not occur in G1 phase, and is regulated in a cell-cycle-dependent manner in human cells. Intriguingly, quantification of both PLA and EdU signals revealed a rough correlation between them (R=0.738). Although PLA signals appeared a little behind EdU during the early S phase (0-4 h), both of them reached the peak at the same time (6 h) followed by a similar decline (Figure 3C). Since EdU incorporation is an indicator of genome replication progress, it strongly argues that cohesin-cohesin interaction occurs in a DNA replication-coupled manner. The time lag of PLA signals compared to EdU levels during the early replication stage is not surprising given that cohesin distributes in an average 67 kb distance along the chromosome in HeLa cells (45).

**Figure 3.**
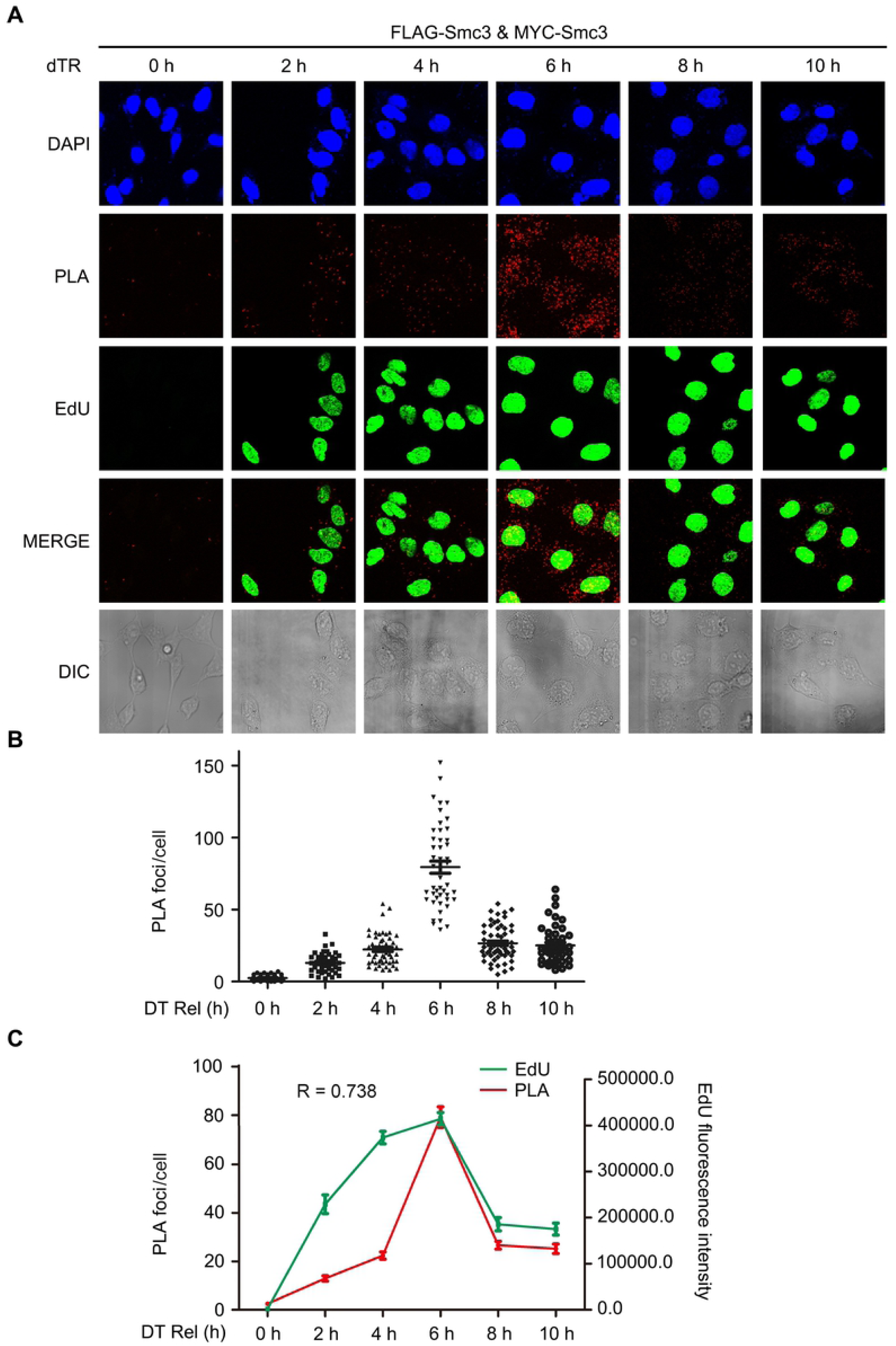
DNA replication-coupled Smc3-Smc3 interaction in human cells. (A) In situ PLA of Smc3-Smc3 in human cells. 293T cells expressing 5FLAG-Smc3 and 13MYC-Smc3 were cultured and synchronized in early S phase by double-thymidine arrest before release (DT Rel) into the fresh media containing EdU. Cells were collected at the indicated time points. PLA was performed by proximity probes against FLAG and MYC. EdU was detected via click-chemistry. (B) Scatter plot of PLA foci per cell throughout the cell cycle. The number of PLA spots within a cell was quantified. At least 50 cells were measured for each time point. (C) Correlation analysis of PLA spots (red) and EdU intensity (green) per cell. The intensity of EdU was measured by ImageJ. The maximal values for in situ PLA and EdU were normalized to allow a comparison between the different assays. Each data point represents an average ± standard error of the mean (SEM) from three biological repeats.

### Intermolecular cohesin interaction is cell-cycle-regulated

To further elucidate how cohesin dimerization is regulated, we investigated it in the synchronized yeast cells. For this purpose, a strain carrying Flag-Scc1 and GFP-Scc1 was grown at 30°C and arrested in G1 by α-factor (0 min). Post-release into S phase, cells were collected at different time points. Then, we carried out GBP-IP of whole-cell extracts. Although Scc1 is expressed in G1 (Figure 4A, lane 3, input/IN), few Scc1 co-precipitated with itself (Figure 4A, upper panel). If we normalized the precipitated GFP-Scc1 (second panel), the co-precipitated 5FLAG-Scc1 gradually increased and peaked around 45 min (S phase) followed by a decline in M phase (Figures 4A, first panel, and 4B). The relative Scc1-Scc1 interaction was quantified as the 5FLAG-Scc1/GFP-Scc1 ratio in the precipitates, which clearly fluctuated with the cell cycle (Figure 4C). Consistent with the results above in human cells, these data suggest that cohesin dimerization occurs exclusively in S phase with a cell-cycle-regulated fashion.

**Figure 4.**
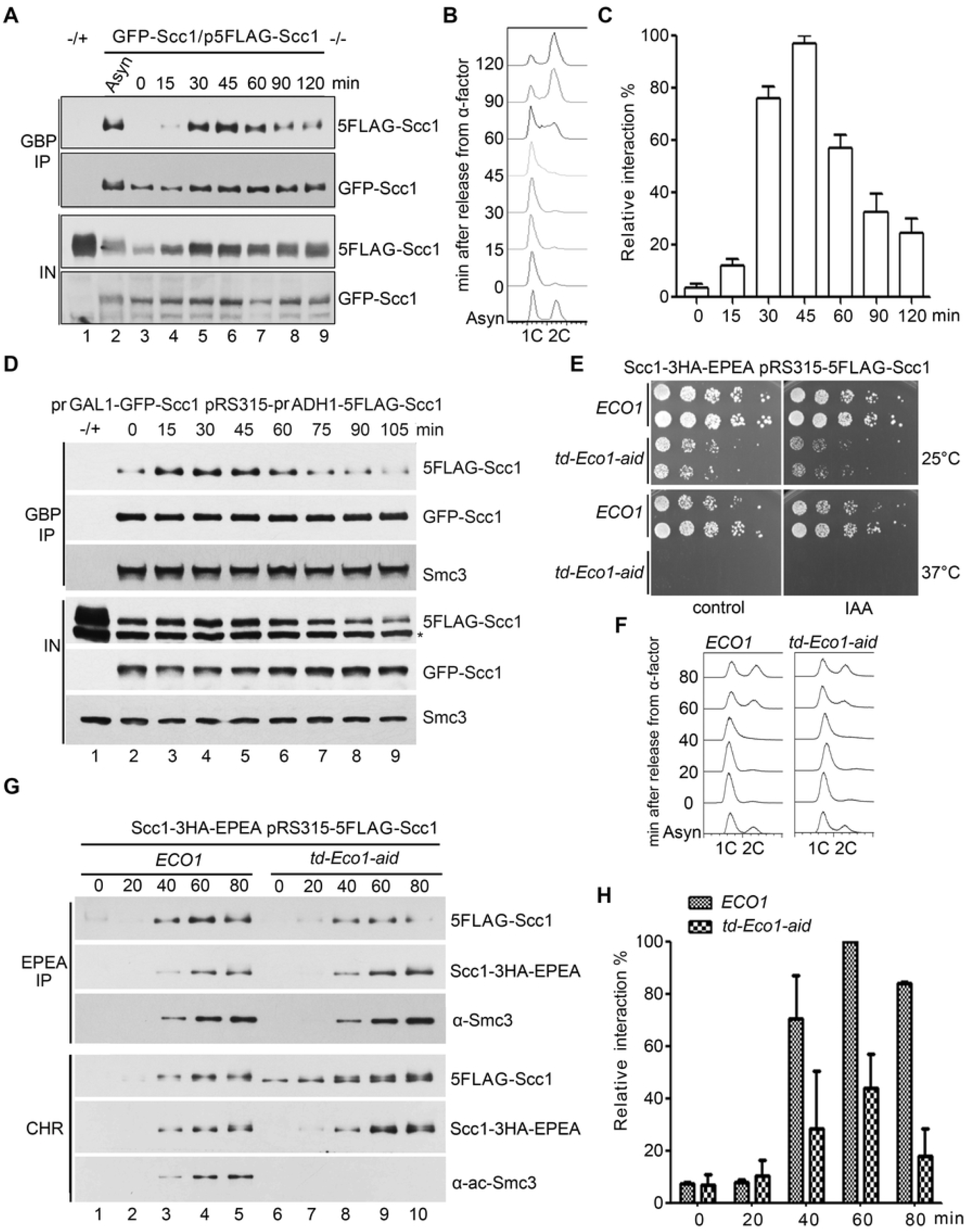
Cohesin-cohesin interaction is regulated by Eco1 during the cell-cycle. (A) The *GFP-SCC1/p5FLAG-SCC1* cells were grown, synchronized in G_1_ by α-factor (0 min) and released into S phase at 25°C for the indicated time. The cell lysates were subjected to GBP-IP and IB against anti-Flag and anti-GFP antibodies. (B) A representative cell cycle profile analyzed by flow cytometry of the samples used in (A). (C) Quantification of the relative intermolecular interaction of cohesin during the cell cycle. The densities of the FLAG-Scc1 and GFP-Scc1 bands in the precipitates were quantified. The ratio of FLAG-Scc1/GFP-Scc1 was calculated to indicate the relative cohesin-cohesin interaction in each sample. The maximum percentage among all samples was normalized to 100%. To ensure the signals were within the linear range, immunoblots with appropriate exposure were quantified by Quantity One (Bio-Rad). (D) Both *GFP-SCC1* and *5FLAG-SCC1* under control of the GAL1 promoter were overexpressed in α-factor arrested cells by galactose. All other experimental conditions were the same as described in (A). (E) Efficient depletion of Eco1 via combined td and aid degrons leads to cell death. The growth of WT (*ECO1*) and Eco1-depletion strains (*td-ECO1-aid*) was examined by spotting on the media with or without IAA at either 25°C or 37°C. (F) Eco1-depletion causes only subtle changes in the cell cycle progression. Representative cell-cycle profiles of WT and Eco1-depletion strains used in (G). After release from G1 arrest, cells were collected at the indicated time at 37°C and analyzed by flow cytometry. (G) Eco1-depletion interferes with cohesin-cohesin interaction on chromatin. Synchronized cells were prepared as in (F). Native chromatin-bound fraction (CHR) was prepared as described in Experimental procedure. Scc1-3HA-EPEA was then precipitated via a C-tag affinity matrix and probed with the indicated antibodies. (H) The relative cohesin-cohesin interaction in the presence or absence of Eco1 was quantified as described in (C).

Notably, there was an increased Scc1 expression during S phase (Figure 4A). To test whether cohesin dimerization in S phase is due to the increased Scc1 protein level, we overexpressed both GFP-Scc1 and 5FLAG-Scc1 by strong promoters. This resulted in a very high level of both versions of Scc1 in G1 (Figure 4D, lane 2, lower panel). However, the Scc1-Scc1 interaction remained very weak at that time and augmented in S phase (Figure 4D, upper panel), similar to that in WT (Figures 4A, upper panel). Meanwhile, the amounts of Smc3 in the precipitates were not significantly changed (Figure 4D, third panel), indicating a constant Scc1-Smc3 interaction throughout the cell cycle. These data suggest that cell-cycle-regulated cohesin dimerization is not merely due to the fluctuation of the Scc1 protein level.

### Cohesin oligomerization shares the common factors as sister chromatid cohesion

The above results suggest a similar cell-cycle-regulated pattern between cohesin-cohesin interaction and sister chromatid cohesion. Notably, both of them occur concomitantly with DNA replication. These facts prompted us to speculate on a functional relationship between the two critical events. To test this notion, we carried out four sets of experiments.

First, we examine whether the vital cohesion establishment factor, Eco1, is required for the cohesin-cohesin interaction. Since Eco1 is essential for cell viability, we combined both temperature-sensitive (td) and auxin-induced (aid) degrons to deplete cellular Eco1 protein. The Ubr1 and Tir1 ubiquitin E3 ligases were induced by galactose. The td and aid degrons were turned on by switching from 25°C to 37°C and adding indole-3-acetic acid (IAA), respectively. These led to cell death (Figure 4E) and abolished S phase Smc3 acetylation (Figure 4G, lanes 7-10, lower panel), indicating effective Eco1 depletion. However, the first S phase progression right after Eco1-depletion was only subtly affected (Figure 4F). Meanwhile, we monitored the intermolecular cohesin interaction during the cell cycle through EPEA-IP and immunoblots in the chromatin-bound fraction (CHR). In WT, the cohesin-cohesin interaction displayed a cell cycle pattern (Figures 4G and 4H) as shown in Figure 4C, but relatively slow which is in accord with the slower cell cycle progression under this condition (Figure 4F). When Eco1 was depleted, co-precipitated 5FLAG-Scc1 was largely decreased (Figures 4G and 4H), whereas the chromatin-associated Scc1 levels of both versions were not much affected (lower panel). Meanwhile, Scc1-Smc3 interaction was not affected either. These data suggest that Eco1 is required for cohesin dimerization, but not for chromatin association of single-rings.

Second, Smc3 acetylation is erased by deacetylase Hos1 in anaphase and recycled in the subsequent cell cycle. So we examined the change of cohesin-cohesin interaction in the absence of Hos1. In the GFP-Scc1/p5FLAG-Scc1 dual-tagged haploid background, WT or mutant cells were cultured and arrested in G2 by nocodazole. Although the amounts of both GFP-Scc1 and 5FLAG-Scc1 were nearly equal in WT and mutant cells (Figure 5A, lanes 5-6), the *hos1*Δ cells showed a significant augment of Scc1-Scc1 interaction (Figure 5A, compare lanes 11 and 12). Similar results were obtained from the diploid cells in which two endogenous Scc1 copies carry the same pair of orthogonal epitopes (Figure 5B, compare lanes 8-10). These results suggest that Hos1 either partially relieves cohesin-cohesin interaction in the M phase or prevents precocious cohesin-cohesin interaction before the S phase.

**Figure 5.**
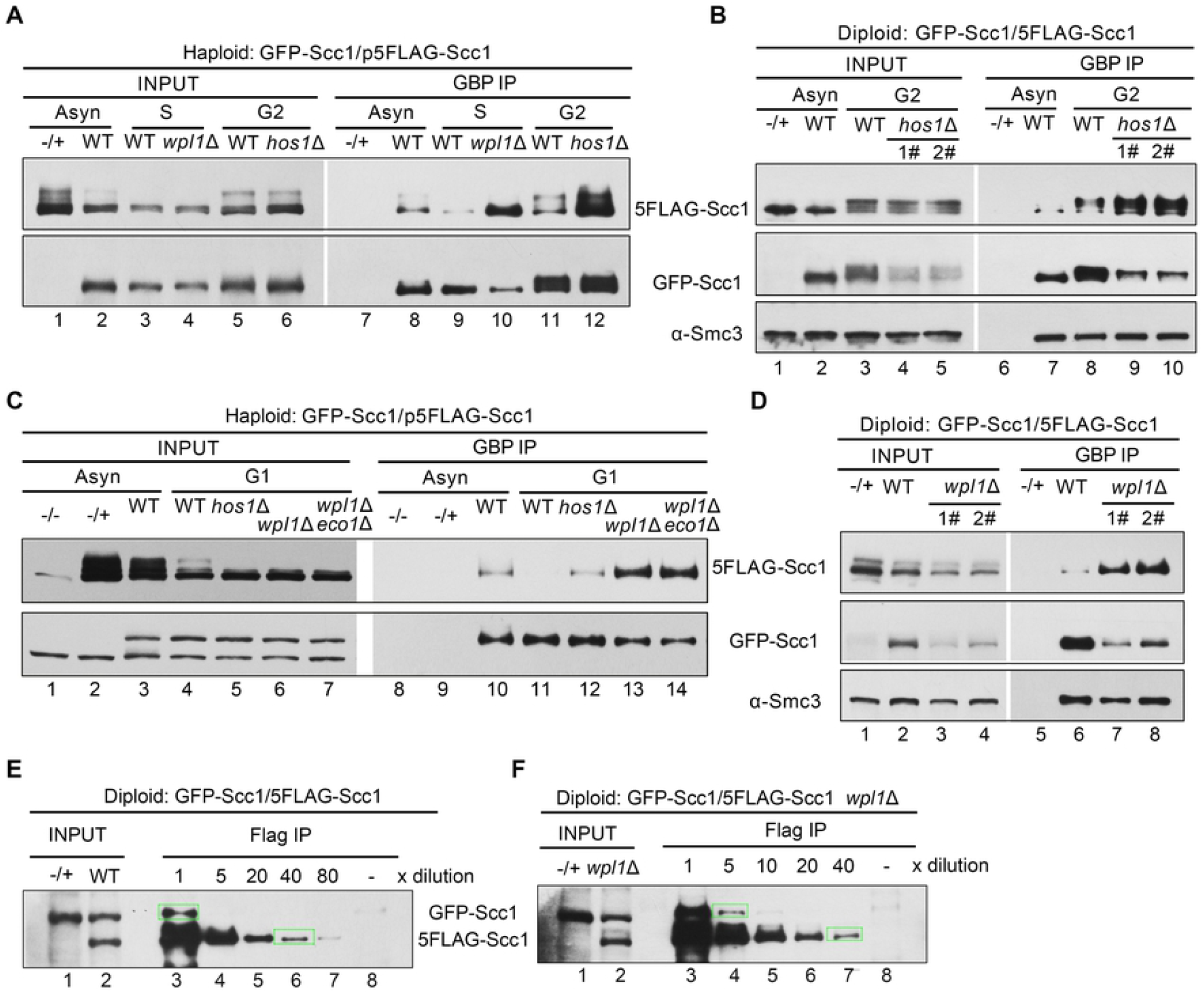
The dimerized cohesin increases once wpl1 or hos1 is depleted. (A) The haploid WT (*GFP-Scc1/p5FLAG-Scc1*), *wpl1*Δ or *hos1*Δ cells were grown with or without synchronization. The S phase cells were obtained by α-factor arrest and release for 60 minutes at 30°C, whereas the G2 cells were arrested by nocodazole. GFP-Scc1 was precipitated via GBP beads from WCE. The proteins were detected via IBs against the indicated antibodies. “-/+” represents the control strain that does not harbor the GFP-tagged version of Scc1. (B) The diploid WT (*GFP-Scc1/5FLAG-Scc1*) or *hos1*Δ cells were grown with or without G2-arrest. The lysates were subjected to GBP-IP and IB as above. 1# and 2# denote the biological repeats. (C) The haploid WT (*GFP-Scc1/p5FLAG-Scc1*), *hos1*Δ, *wpl1*Δ, or *wpl1*Δ*eco1*Δ cells were grown with or without G1-arrest. GBP-IPs and IBs were performed as above. (D) The diploid WT (*GFP-Scc1/5FLAG-Scc1*) or *wpl1*Δ cells were grown exponentially. The lysates were subjected to GBP-IP and IB. 1# and 2# denote the biological repeats. (E, F) The diploid WT (*GFP-Scc1/5FLAG-Scc1*) (E) or *wpl1*Δ (F) cells were grown exponentially. The lysates were precipitated by anti-FLAG M2 beads. A series of dilutions (5×, 10×, 20×, 40×, 80×) of the samples were probed by anti-Scc1 antibodies. The indicated relative density of the band in a rectangular marquee was measured by Quantity One (BioRad). The percentage of cohesin dimers was calculated as described in Experimental Procedures. “-/+” represents the control strain that does not harbor the 5FLAG-tagged version of Scc1.

Third, prior to Eco1-dependent cohesion establishment, cohesin remains dynamic on chromatin due to the destabilized activity of Wpl1. The essential function of *ECO1* can be bypassed by *WPL1* deletion (33, 34). So, we next compared the cohesin-cohesin interaction in the presence or absence of Wpl1. The experiments were basically conducted as described for *hos1*Δ. When *WPL1* was deleted, cohesin oligomerization increased prominently in either G1 (Figure 5C, compare lanes 11 and 13) or S (Figure 5A, compare lanes 9 and 10). Consistently, in asynchronized diploid cells, a dramatic increase was observed in the absence of Wpl1 as well (Figure 5D, compare lanes 6-8). These data indicate that Wpl1 prevents the cohesin-cohesin interaction, correlating with a loose and dynamic association of cohesin with chromatin in G1.

Fourth, based on the ratio of GFP-Scc1/5FLAG-Scc1 in the cell extracts and precipitates, we estimated the percentage of cohesin dimers at the endogenous protein levels in diploid cells using a method published recently (41). The cellular levels of GFP-Scc1 and 5FLAG-Scc1 were measured nearly identical (Figure 5E). Through serial dilutions of the precipitates, we quantified the band densities of GFP-Scc1 and 5FLAG-Scc1 probed by anti-Scc1 antibodies in the same gel. The percentage of cohesin dimers was roughly estimated through the following formula:

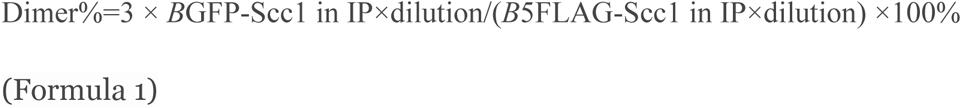

In asynchronized WT cells, ∼20% of cohesins could be detected in the dimeric state (Figure 5E). Intriguingly, ∼20%-30% of Smc3 is acetylated in budding yeast in a previous report (35). Using a similar method, Cattoglio et al recently reported that cohesin dimers occupy at least ∼8% in mESCs (43). When *WPL1* was deleted, cohesin dimers increased up to 40% in yeast (Figure 5F). This result suggests that Wpl1, the anti-establishment factor of cohesion, acts as a negative regulator in cohesin dimerization as well. Taken together, all these lines of evidence support that cohesin dimerization is cell-cycle-regulated as the sister chromatid cohesion cycle by the same mechanisms (i.e., Wpl1/Eco1/Hos1).

Here, we show that cohesin is dimerized in S phase and monomerized again in mitotis and G1, which is controlled by common regulators (Eco1, Wpl1, Hos1) as the sister chromatid cohesion/dissolution cycle. Besides these biochemical evidence described here and literature (20, 43), genetic interactions also support cohesin-cohesin interactions (19). Both yeast cohesin and prokaryotic SMC condensin have been proposed to act as dimers in extruding DNA loops (48, 49). Therefore, besides the canonical single-ring structure, dimerization or oligomerization may provide an additional mechanism for cohesin to execute various functions in sister chromatid cohesion, DNA repair, chromatin loop extrusion, high-order chromatin organization (21).

## Materials and Methods

### Strain and plasmid construction

Strains and plasmids used in this study are listed in Tables S1 and S2, respectively.

### Preparation of antibodies

To raise polyclonal antibodies specific to Scc1 and Smc3, purified Scc1N (1-333 a.a.) and Smc3 hinge domain were used to immunize rabbits. Polyclonal antibodies were affinity purified. Scc1 and Smc3 beads were prepared by immobilizing purified Scc1N and Smc3 proteins to NHS-activated agarose beads as recommended by the manufacturer (GE Healthcare).

### Cell synchronization and flow cytometry analysis

Cells were grown to logarithmic phase, 7.5 μg/ml of α-factor was added for cell synchronization in G1-phase. After washing twice, G1 arrested cells were released in fresh medium and continued growth for the indicated time. Cells were collected and fixed with 70% ethanol and then processed for flow cytometry using a BECKMAN Cytoflex-S.

### Conditional depletion of cellular Eco1 protein

The efficient depletion of endogenous Eco1 protein was achieved through a two-degron strategy. Temperature-inducible (td) and auxin-inducible (aid) degrons were added to the N and C terminus of Eco1, respectively. The corresponding two ubiquitin ligases (E3), UBR1 and OsTIR1, were integrated into the genomic *UBR1* locus under control of the galactose-inducible Gal1-10 promoter. Cells were first grown at 25°C in rich medium contains 0.1 mM Cu^2+^ supplemented with 2% raffinose before transferring to 2% galactose to induce the expression of two E3s. Two degrons were turned on by adding 1 mM indole-3-acetic acid (IAA) (for aid) and switching to 37°C (for td) for 2 hr. The protein level of Eco1-MYC was measure by IB with anti-MYC and anti-Smc3ac antibodies.

### Whole-cell extracts (WCE) and immunoblotting (IB)

WCE of one hundred OD600 units of asynchronized or synchronized cells were prepared by glass bead beating (Mini-Beadbeater-16, Biospec,USA) in lysis buffer {(50 mM HEPES/KOH pH7.4, 150 mM NaCl, 1 mM EDTA, 10% Glycerol, 1 mM DTT, 1 mM PMSF, Protease inhibitor tablets (EDTA free, Roche)}. Protein samples were separated by SDS-polyacrylamide gel electrophoresis (SDS-PAGE) and immunoblotted with the antibodies specifically indicated in each figure. Antibodies used in this study are as follows: mouse anti-FLAG M2-specific monoclonal antibody (1:1000, Sigma), rabbit polyclonal anti-GFP (1:500, GeneScript), mouse anti-HA 16B12 (1:1000, Millipore), anti-ac-Smc3, polyclonal anti-Smc3 (1:1000), polyclonal anti-Scc1 (1:1000). HRP-conjugated anti-rabbit or anti-mouse IgG was used as the secondary antibody (1:10000, Sigma).

### Immunoprecipitation (IP)

Monoclonal GBP agarose, monoclonal anti-EPEA agarose (Thermo Fisher) and monoclonal anti-Flag M2 affinity gel (Sigma-Aldrich) were used for IP. IP was performed using strains co-expressing the tagged versions of each protein as indicated in each figure. After three washes, the proteins specifically associated with beads were boiled and analyzed by SDS-PAGE and IB using the indicated antibodies.

### Glycerol gradient centrifugation

The native protein complexes in the peptide eluates after FLAG-IPs were concentrated and applied to the top of a 10–30% glycerol gradient in EBX-3 buffer {50 mM HEPES/KOH pH7.5, 150 mM KCl, 2.5 mM MgOAc, 0.1 mM ZnOAc, 2 mM NaF, 0.5 mM spermidine, 20 mM Glycerophosphate, 1 mM ATP, 1 mM DTT, 1 mM PMSF, Protease inhibitor tablets (EDTA free, Roche)}. The gradients were centrifuged in a P55ST2 swinging bucket rotor (Hitachi CP100NX ultracentrifuge) at 120,000g for 9 h using slow deceleration. After centrifugation, the fractions were collected from the top of the gradient and subjected to SDS-PAGE and immunoblots. Aldlase (158 kDa) and thyroglobulin (669 kDa) were used as size markers.

### CXMS (Cross-Linking Mass Spectrometry)

5FLAG-Scc3 was prepared by FLAG-IP of yeast WCE and FLAG peptide elution. About 15 μg of purified Scc3 in a volume of 15 μl was cross-linked through incubation with the lysine cross-linker disuccinimidyl suberate (DSS) at a final concentration of 0.5 mM for 1 h at room temperature. The final concentration of 20 mM NH_4_HCO_3_ was added to quench the reaction. The cross-linked proteins were precipitated with ice-cold acetone of 4-5 fold volume at −20°C overnight, resuspended in 8 M urea, 100 mM Tris, pH 8.5. After trypsin digestion, the LC-MS/MS analysis was performed on an Easy-nLC 1000 UHPLC (Thermo Fisher Scientific) coupled to a Q Exactive HF Orbitrap mass spectrometer (Thermo Fisher Scientific). Peptides were loaded on a pre-column (75 μm inner diameter, 4 cm long, packed with ODS-AQ 12 nm–10 mm beads from YMC Co., Ltd.) and separated on an analytical column (75 μm inner diameter, 13 cm long, packed with ReproSil-Pur C18-AQ 1.9 μm 120 A° resin from Dr. Maisch GmbH) using an linear gradient of 0–35% buffer B (100% acetonitrile and 0.1% formic acid) at a flow rate of 250 nl/min over 73 min. The top 20 most intense precursor ions from each full scan (resolution 60,000) were isolated for HCD MS2 (resolution 15,000; NCE 27) with a dynamic exclusion time of 30 s. Precursors with 1+ or unassigned charge states were excluded. pLink was used to identify cross-linked peptides with the cutoffs of FDR, 5% and E_value,0.001.

### Disulfide crosslinking to capture site-specific protein-protein interactions in vivo

The Scc3-Scc3 interaction was captured by a disulfide crosslinking method in yeast cells (44). The indicated pairs of amino acid residues in Scc3 were substituted by cysteine. WT or mutant cells were cultured in 5 mL of CSM media (without cysteine) at 30°C to OD_600_ of 0.5 before the addition of 180 μM 4,4’-dipyridyl disulfide (4-DPS, Sigma Aldrich). The cultures were resumed for 20 min and then quenched with 20% trichloroacetic acid (TCA). The cells were pelleted and washed with 20% TCA before homogenization in the presence of 400 μL 20% TCA and ∼450 μL of glass beads using Mini-Beadbeater-16 (Biospec, USA). N-ethyl maleimide (NEM) was added to prevent any free thiol groups from crosslinking after cell lysis. Proteins were extracted and separated by non-reducing and reducing sodium dodecyl sulfate-polyacrylamide gel electrophoresis (SDS-PAGE) for immunoblots against the indicated antibodies as previously described (44).

### Proximity ligation assay (PLA)

The PLA was performed as described previously (50). Briefly, HeLa cells were fixed with 4% paraformaldehyde (Sigma-Aldrich, USA) in PBS for 15 min, permeabilized with 0.1% Triton X-100 (Sigma) for 5 min, and blocked for 1 h with a blocking solution(250 μg/ml BSA, 2.5 μg/ml Sonicated salmon sperm DNA, 5 mM EDTA, 0.05% Tween 20 in PBS). Cells were washed with PBS and incubated in two primary antibodies, the primary antibodies used were as follows: mouse monoclonal anti-FLAG (1:100; Sigma), rabbit polyclonal anti-MYC (1:100; 16286-1-AP, proteintech). After washing with PBS, samples were incubated with secondary antibodies conjugated with PLA probes for 1 h at 37°C. After washing with PBS, samples were incubated with ligation–ligase solution for 30 min at 37°C. Then the samples were washed with PBS and continued with amplification-polymerase solution incubation for 90 min at 37°C. Add the detection mix and incubate for 30 min at 37°C. From now on keep the slide in the dark. After washing three times for 5 min each with PBS, slides were mounted using Duolink in situ Mounting Medium with DAPI. Pictures were taken using a fluorescent microscope (Leica DIM8).

### Native chromatin fractionation

Native chromatin fraction was performed as described (51, 52) with minor modifications. Yeast cells of 200 OD600 units were spheroplasted by 75 U/ml lyticase. Crude extracts were prepared by Triton X-100 treatment and fractionated via sucrose cushion in 500 μl of EBX-3 buffer {50 mM HEPES/KOH pH7.4, 150 mM NaCl, 2.5 mM MgCl_2_, 0.1 mM ZnOAc, 5 mM NaF, 1 mM NaVO_4_, 10 mM β-Glycerophosphate, 1 mM ATP, 1 mM DTT, 1 mM PMSF, Protease inhibitor tablets (EDTA free, Roche)}. The supernatant contains non-chromatin bound proteins. Chromatin-bound proteins (CHR) in the pellet were released by incubation in EBX-3 buffer containing 500 U/ml of Benzonase (Sigma) for 60 min at 4°C.

## Acknowledgments

We thank Dr. Katsuhiko Shirahige (Tokyo Institute of Technology) for anti-Smc3ac antibodies; and members of the Lou lab for helpful discussion and comments on the manuscript.

This work was supported by the National Natural Science Foundation of China 31630005, 31770084 and 31771382; the National Basic Research Program (973 Program) of China (2014CB849801); Program for Extramural Scientists of the State Key Laboratory of Agrobiotechnology 2018SKLAB6-5.

---

**Table S1.** Plasmids used in this study

| Plasmid | Base plasmid/Genotype | Source |
| --- | --- | --- |
| pRS315-5Flag-SCC1 | *amp<sup>r</sup>/LEU2 5Flag-SCC1* | This study |
| pPADH1-5Flag-SCC1 | *amp<sup>r</sup>/HIS3 5Flag-SCC1* | This study |
| pPADH1-5Flag-SCC3 | *amp<sup>r</sup>/LEU2 5Flag-SCC3* | This study |
| pPADH1-13MYC-SCC3 | *amp<sup>r</sup>/HIS3 13MYC-SCC3* | This study |
| pPADH1-5Flag- scc3 K99C | *amp<sup>r</sup>/HIS3 5Flag- scc3 K99C* | This study |
| pPADH1-5Flag- scc3 K764C | *amp<sup>r</sup>/HIS3 5Flag- scc3 K764C* | This study |
| pPADH1-13MYC-scc3 K1076C | *amp<sup>r</sup>/LEU2 13MYC-scc3 K1076C* | This study |
| pPADH1-13MYC- scc3 K1057C | *amp<sup>r</sup> /LEU2 13MYC- scc3 K1057C* | This study |
| pRS313-ECO1 | *amp<sup>r</sup> /HIS3 ECO1* | This study |
| pRS313-PCUP1-td-ECO1-13MYC-aid | *amp<sup>r</sup> /HIS3 td-ECO1-13MYC-aid* | This study |
| pRK5-FLAG-SMC3 | *amp<sup>r</sup>/FLAG-SMC3* | This study |
| pRK5-MYC-SMC3 | *amp<sup>r</sup>/MYC-SMC3* | This study |

**Table S2.**
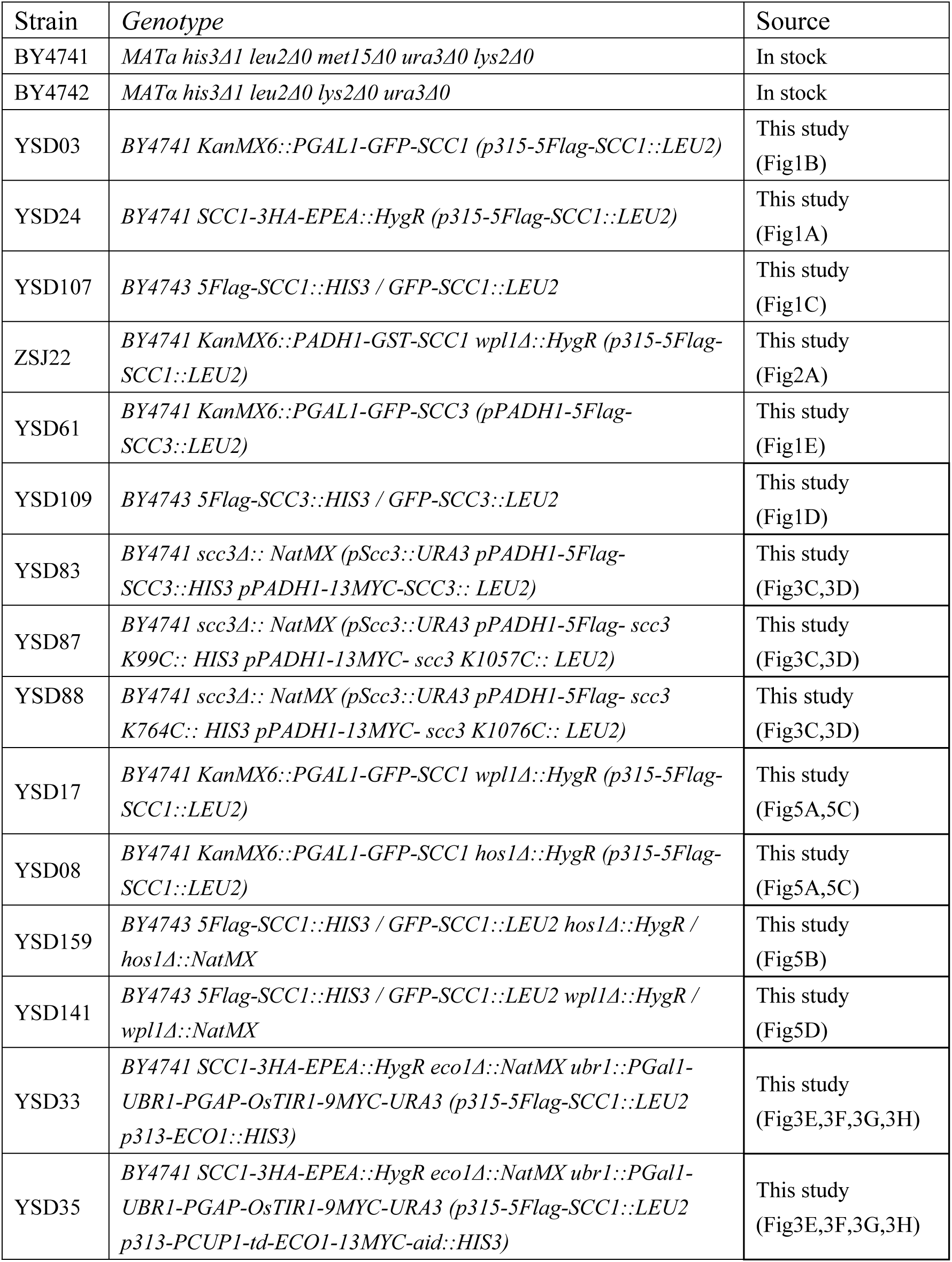
Strains used in this study.

## References

1. Onn I, Heidinger-Pauli JM, Guacci V, Unal E, Koshland DE. Sister chromatid cohesion: a simple concept with a complex reality. Annu Rev Cell Dev Biol. 2008;24:105–29.

2. Peters J-M, Tedeschi A, Schmitz J. The cohesin complex and its roles in chromosome biology. Genes & Development. 2008;22(22):3089–114.

3. Nasmyth K, Haering CH. Cohesin: its roles and mechanisms. Annu Rev Genet. 2009;43:525–58.

4. Jeppsson K, Kanno T, Shirahige K, Sjogren C. The maintenance of chromosome structure: positioning and functioning of SMC complexes. Nat Rev Mol Cell Biol. 2014;15(9):601–14.

5. Uhlmann F. SMC complexes: from DNA to chromosomes. Nature Reviews Molecular Cell Biology. 2016;17:399.

6. Nasmyth K. Cohesin: a catenase with separate entry and exit gates? Nature Cell Biology. 2011;13(10):1170–7.

7. Gligoris T, Lowe J. Structural Insights into Ring Formation of Cohesin and Related Smc Complexes. Trends in cell biology. 2016;26(9):680–93.

8. Li Y, Muir KW, Bowler MW, Metz J, Haering CH, Panne D. Structural basis for Scc3-dependent cohesin recruitment to chromatin. Elife. 2018;7.

9. Haering CH, Löwe J, Hochwagen A, Nasmyth K. Molecular Architecture of SMC Proteins and the Yeast Cohesin Complex. Molecular Cell. 2002;9(4):773–88.

10. Holzmann J, Fuchs J, Pichler P, Peters JM, Mechtler K. Lesson from the stoichiometry determination of the cohesin complex: a short protease mediated elution increases the recovery from cross-linked antibody-conjugated beads. J Proteome Res. 2011;10(2):780–9.

11. Holzmann J, Politi AZ, Nagasaka K, Hantsche-Grininger M, Walther N, Koch B, et al. Absolute quantification of cohesin, CTCF and their regulators in human cells. Elife. 2019;8.

12. Gruber S, Haering CH, Nasmyth K. Chromosomal cohesin forms a ring. Cell. 2003;112(6):765–77.

13. Ivanov D, Nasmyth K. A topological interaction between cohesin rings and a circular minichromosome. Cell. 2005;122(6):849–60.

14. Chan KL, Roig MB, Hu B, Beckouet F, Metson J, Nasmyth K. Cohesin’s DNA exit gate is distinct from its entrance gate and is regulated by acetylation. Cell. 2012;150(5):961–74.

15. Murayama Y, Uhlmann F. Biochemical reconstitution of topological DNA binding by the cohesin ring. Nature. 2014;505(7483):367–71.

16. Huis in ‘t Veld PJ, Herzog F, Ladurner R, Davidson IF, Piric S, Kreidl E, et al. Characterization of a DNA exit gate in the human cohesin ring. Science. 2014;346(6212):968–72.

17. Srinivasan M, Scheinost JC, Petela NJ, Gligoris TG, Wissler M, Ogushi S, et al. The Cohesin Ring Uses Its Hinge to Organize DNA Using Non-topological as well as Topological Mechanisms. Cell. 2018;173(6):1508-19.e18.

18. Murayama Y, Samora CP, Kurokawa Y, Iwasaki H, Uhlmann F. Establishment of DNA-DNA Interactions by the Cohesin Ring. Cell. 2018;172(3):465-77.e15.

19. Eng T, Guacci V, Koshland D. Interallelic complementation provides functional evidence for cohesin-cohesin interactions on DNA. Molecular biology of the cell. 2015;26(23):4224–35.

20. Zhang N, Kuznetsov SG, Sharan SK, Li K, Rao PH, Pati D. A handcuff model for the cohesin complex. The Journal of cell biology. 2008;183(6):1019–31.

21. Skibbens RV. Of Rings and Rods: Regulating Cohesin Entrapment of DNA to Generate Intra- and Intermolecular Tethers. PLoS Genet. 2016;12(10):e1006337.

22. Hansen AS, Pustova I, Cattoglio C, Tjian R, Darzacq X. CTCF and cohesin regulate chromatin loop stability with distinct dynamics. eLife. 2017;6:e25776.

23. Hirano T, Nishiyama T, Shirahige K. Hot debate in hot springs: Report on the second international meeting on SMC proteins. Genes to Cells. 2017;22(11):934–8.

24. Peters J-M, Nishiyama T. Sister Chromatid Cohesion. Cold Spring Harbor Perspectives in Biology. 2012;4(11).

25. Morales C, Losada A. Establishing and dissolving cohesion during the vertebrate cell cycle. Current Opinion in Cell Biology. 2018;52:51–7.

26. Ciosk R, Shirayama M, Shevchenko A, Tanaka T, Toth A, Shevchenko A, et al. Cohesin’s Binding to Chromosomes Depends on a Separate Complex Consisting of Scc2 and Scc4 Proteins. Molecular Cell. 2000;5(2):243–54.

27. Sherwood R, Takahashi TS, Jallepalli PV. Sister acts: coordinating DNA replication and cohesion establishment. Genes & Development. 2010;24(24):2723–31.

28. Makrantoni V, Marston AL. Cohesin and chromosome segregation. Current Biology. 2018;28(12):R688–R93.

29. Skibbens RV, Corson LB, Koshland D, Hieter P. Ctf7p is essential for sister chromatid cohesion and links mitotic chromosome structure to the DNA replication machinery. Genes & Development. 1999;13(3):307–19.

30. Toth A, Ciosk R, Uhlmann F, Galova M, Schleiffer A, Nasmyth K. Yeast cohesin complex requires a conserved protein, Eco1p(Ctf7), to establish cohesion between sister chromatids during DNA replication. Genes Dev. 1999;13(3):320–33.

31. Skibbens RV. Establishment of Sister Chromatid Cohesion. Current Biology. 2009;19(24):R1126–R32.

32. Uhlmann F. A matter of choice: the establishment of sister chromatid cohesion. EMBO reports. 2009;10(10):1095–102.

33. Rolef Ben-Shahar T, Heeger S, Lehane C, East P, Flynn H, Skehel M, et al. Eco1-dependent cohesin acetylation during establishment of sister chromatid cohesion. Science. 2008;321(5888):563–6.

34. Unal E, Heidinger-Pauli JM, Kim W, Guacci V, Onn I, Gygi SP, et al. A molecular determinant for the establishment of sister chromatid cohesion. Science. 2008;321(5888):566–9.

35. Zhang J, Shi X, Li Y, Kim B-J, Jia J, Huang Z, et al. Acetylation of Smc3 by Eco1 Is Required for S Phase Sister Chromatid Cohesion in Both Human and Yeast. Molecular Cell. 2008;31(1):143–51.

36. Sutani T, Kawaguchi T, Kanno R, Itoh T, Shirahige K. Budding yeast Wpl1(Rad61)-Pds5 complex counteracts sister chromatid cohesion-establishing reaction. Curr Biol. 2009;19(6):492–7.

37. Tanaka K, Yonekawa T, Kawasaki Y, Kai M, Furuya K, Iwasaki M, et al. Fission yeast Eso1p is required for establishing sister chromatid cohesion during S phase. Mol Cell Biol. 2000;20(10):3459–69.

38. Feytout A, Vaur S, Genier S, Vazquez S, Javerzat JP. Psm3 acetylation on conserved lysine residues is dispensable for viability in fission yeast but contributes to Eso1-mediated sister chromatid cohesion by antagonizing Wpl1. Mol Cell Biol. 2011;31(8):1771–86.

39. Hauf S, Roitinger E, Koch B, Dittrich CM, Mechtler K, Peters JM. Dissociation of cohesin from chromosome arms and loss of arm cohesion during early mitosis depends on phosphorylation of SA2. PLoS biology. 2005;3(3):e69.

40. Quan Y, Xia Y, Liu L, Cui J, Li Z, Cao Q, et al. Cell-Cycle-Regulated Interaction between Mcm10 and Double Hexameric Mcm2-7 Is Required for Helicase Splitting and Activation during S Phase. Cell reports. 2015;13(11):2576–86.

41. Liu L, Zhang Y, Zhang J, Wang J-H, Cao Q, Li Z, et al. Characterization of the dimeric CMG/pre-initiation complex and its transition into DNA replication forks. Cellular and Molecular Life Sciences. 2019.

42. Maradeo ME, Skibbens RV. Epitope tag-induced synthetic lethality between cohesin subunits and Ctf7/Eco1 acetyltransferase. FEBS letters. 2010;584(18):4037–40.

43. Cattoglio C, Pustova I, Walther N, Ho JJ, Hantsche-Grininger M, Inouye CJ, et al. Determining cellular CTCF and cohesin abundances to constrain 3D genome models. eLife. 2019;8:e40164.

44. Mohan C, Kim LM, Hollar N, Li T, Paulissen E, Leung CT, et al. VivosX, a disulfide crosslinking method to capture site-specific, protein-protein interactions in yeast and human cells. eLife. 2018;7:e36654.

45. Holzmann J, Politi AZ, Nagasaka K, Hantsche-Grininger M, Walther N, Koch B, et al. Absolute quantification of cohesin, CTCF and their regulators in human cells. eLife. 2019;8:e46269.

46. Uhlmann F, Nasmyth K. Cohesion between sister chromatids must be established during DNA replication. Curr Biol. 1998;8(20):1095–101.

47. Soderberg O, Leuchowius KJ, Gullberg M, Jarvius M, Weibrecht I, Larsson LG, et al. Characterizing proteins and their interactions in cells and tissues using the in situ proximity ligation assay. Methods. 2008;45(3):227–32.

48. Kim Y, Shi Z, Zhang H, Finkelstein IJ, Yu H. Human cohesin compacts DNA by loop extrusion. Science. 2019;366(6471):1345–9.

49. Wang X, Brandão HB, Le TBK, Laub MT, Rudner DZ. Bacillus subtilis SMC complexes juxtapose chromosome arms as they travel from origin to terminus. Science (New York, NY). 2017;355(6324):524–7.

50. Weibrecht I, Leuchowius KJ, Clausson CM, Conze T, Jarvius M, Howell WM, et al. Proximity ligation assays: a recent addition to the proteomics toolbox. Expert Rev Proteomics. 2010;7(3):401–9.

51. Sheu YJ, Stillman B. Cdc7-Dbf4 phosphorylates MCM proteins via a docking site-mediated mechanism to promote S phase progression. Mol Cell. 2006;24(1):101–13.

52. Quan Y, Xia Y, Liu L, Cui J, Li Z, Cao Q, et al. Cell-Cycle-Regulated Interaction between Mcm10 and Double Hexameric Mcm2-7 Is Required for Helicase Splitting and Activation during S Phase. Cell reports. 2015;13(11):2576–86.

